# Lysophosphatidic Acid (LPA) Salivary Species Detection and Whole-mount LPA Receptor Localization in Mouse Salivary Gland

**DOI:** 10.64898/2026.04.28.721492

**Authors:** D. Roselyn Cerutis, Devendra Kumar, Michael G. Nichols, Gabrielle Rae Roemer, Maxwell Emmett Fluent, Takanari Miyamoto, Yazen Alnouti

**Author notes:** Other authors.

## Abstract

This study builds on our previous findings on the role of salivary lysophosphatidic acid (LPA) species in humans to investigate their presence, together with salivary gland LPA receptor (LPAR) expression in a *Porphyromonas gingivalis*-infected murine (C57BL/6J) model of periodontal disease (PD). Utilizing LC-MS/MS for LPA analysis alongside confocal LPAR imaging and second harmonic (SHG) imaging for collagen visualization, we compared mouse salivary LPA levels and gland LPAR expression to previously established human and mouse data. The findings reveal that while healthy mouse saliva maintains low homeostatic LPA levels, PD triggers an ∼ 10-fold increase, mirroring the elevation we observed in PD patients. Furthermore, the study confirmed the presence of LPA1, LPA3, and LPA4 within submandibular gland (SMG) tissue. Notably, LPA3 was identified as the most widely distributed subtype, while providing the first evidence of LPA4 expression in adult mouse salivary glands. The presence of multiple LPARs suggests that LPA signaling is a critical factor in salivary gland biology. The documented existence of multiple LPARs within salivary glands indicates that they must be taken into consideration in future research concerning autoimmune conditions, and in pharmacological studies involving drugs that impact salivary gland biology and secretory function.

## Introduction

Lysophosphatidic acid (LPA) is a multipotent small lipid mediator that plays key roles in development, homeostasis and in pathophysiology. It is found at low levels in most normal bodily fluids, but in pathology, it increases to pharmacologic levels and often contributes to the disease processes in many systems [reviewed in 1; 2].

Sugiura et al. (2002) [3] first showed that healthy human whole saliva contains multiple LPA species, the major ones being LPA16:0, LPA18:0, and LPA18:1. The oral mucosa’s lysophospholipase D converts salivary gland-secreted lysophosphatidylcholine (LPC, the first direct precursor of LPA) to LPA; another enzyme, glycerophosphodiesterase 7, is a newly proposed candidate to also produce LPA in mixed saliva [4].

We determined that these salivary LPA species preferentially control the regenerative responses of human gingival (GF) and periodontal ligament fibroblasts (PDLF) [5,6], and that the Ca^2+^ signaling responses of GF and PDLF are preferentially controlled through the LPA receptor (LPAR) subtypes LPA1 and LPA3, with negligible -if any - contribution from LPA2. [7]. We also found that LPA species levels in human saliva and gingival crevicular fluid (GCF) increase with the severity of periodontal disease (PD) [8]. Multiple LPAR subtypes have been shown to be involved in the modulation of human endothelial integrity and inflammation [9]. LPA 18:1 significantly modulates a multitude of inflammatory transcripts in human oral fibroblasts [10].

The submandibular gland (SMG) is a major salivary gland located beneath the mandible in both humans and mice. In mice (and rats) the SMG is the largest salivary gland [11]. Structurally, it has three main components in its connective tissue: acini, ducts, and myoepithelial cells. Acini contain serous cells and mucous cells. Serous cells produce a watery, enzyme-rich secretion, while mucous cells produce a thicker, mucin-rich secretion. Ducts carry the secretions produced by the acini to the oral cavity. The SMG has two duct types: intercalated ducts and striated ducts. The small intercalated ducts connect the acini to the larger, more complex striated ducts which modify saliva composition by reabsorbing ions and secreting bicarbonate. Contractile myoepithelial cells surround the acini and ducts to squeeze the secretions out of the acini and into the ducts.

LPA is essential to regulating many key aspects of oral physiology and pathophysiology, starting with development. Noguchi et al. (2006) [12] determined that LPA and epidermal growth factor (EGF) cooperate to induce branching morphogenesis of embryonic mouse salivary submandibular gland (eSMG) epithelium. Using stearyl-LPA, a non-hydrolyzable LPA analog, they showed that the analog plus EGF were sufficient to stimulate branching morphogenesis. They separated the eSMG into epithelial and mesenchymal parts and used reverse transcriptase-polymerase chain reaction (RT-PCR) to investigate the presence of *Lpa1, Lpa2*, and *Lpa3* mRNA. *Lpa1* and *Lpa2* showed equal expression in the epithelial and mesenchymal eSMG, *Lpa3* was abundant in the epithelial part, and *Lpa4* was expressed higher in the mesenchymal eSMG.

More lines of evidence implicate the importance of LPA to oral biology: it controls essential elements-bone biology, epithelial barrier integrity, and inflammation [reviewed in 13,14, 15, 16]. Further, the Na-K-Cl co-transporter NKCC1 is a critical component for saliva production. Basolateral NKCC1 play a critical role in fluid secretion by promoting the intracellular accumulation of Cl^−^ above its equilibrium potential [17,18] and is the main mechanism for acinar cells concentrating Cl^−^[19]. Significantly, research on related secretory tissues (like the choroid plexus) suggests that LPA can act as an agonist that activates NKCC1 to drive fluid secretion [20] via acting as an agonist of the choroidal transient receptor potential vanilloid 4 (TRPV4) channel [21]. Although this possibility (to our knowledge) has not been investigated for salivary glands, we propose that LPA is likely to play the same critical role regulating NKCC1for saliva production.

Normal salivary gland secretion can be affected/diminished by some infections (bacterial and viral); systemic conditions like diabetes, Sjögren’s syndrome (SS), and neurological diseases; and a wide array of medications, including anti-hypertensives, β-blockers, antidepressants, and antihistamines; the salivary glands also often sustain off-target damage from radiation treatments for head or neck cancers. [22,23,24].

SS is an inflammatory condition where salivary gland epithelial cells are one of the main targets of the autoimmune attack and so develop secretory hypofunction [reviewed in 25]. Dysfunction of salivary glands is reflected in oral-systemic inflammatory pathways, which produce mediators that negatively influence chronic conditions like cardiovascular disease, diabetes mellitus, arthritis, and others [26, 23]. LPA contributes to salivary gland pathophysiology in SS. In preclinical studies, the LPAR1/LPAR3 antagonist Ki16425 was found to restore saliva and tear secretion via reducing the serum and lacrimal gland interleukin-17 [(IL-17) elevation that LPA induces in SS through the ROCK2 and p38 MAPK pathways] [27,28].

Iperi et al. (2025) [29] used transcriptomic, methylomic, and metabolomic SS patient datasets from the PRECISESADS study to further implicate LPA in SS pathology. They found that lysophosphatidylcholine (LysoPC) species LysoPC (22:6) and LysoPC (22:5), the direct substrates for autotaxin (the main LPA-generating enzyme) were elevated. They also identified that LPAR6 is expressed on B lymphocytes -and would bind that LPA-and so proposed making the LPA–LPAR6 axis a candidate for therapeutic modulation of SS.

Collagen is a major extracellular matrix (ECM) component that provides structural integrity and mechanical strength to bones, organs, skin, and glands. The way collagen type I is structurally arranged within tissues plays a major role in determining their architectural and biomechanical properties, which are closely linked to tissue physiology and function.

During inflammation, collagen undergoes detrimental changes, including degradation [30, 31,32]. Collagen fibrils crosslink to form fibrillar networks that support tissue architecture [33]. Because the structure of organs and glands depends heavily on this collagen-based support, maintaining collagen structural integrity is essential for healthy function.

Second harmonic generation (SHG) microscopy enables high-resolution, highly detailed, label-free visualization of collagen fiber density and fibril organization [reviewed in 34;35], making it a powerful tool for assessing collagen structure within tissues. This ability for non-invasive visualization is a boon to acquire a better understanding of how collagen density and organization is impacted by treatments and pathological conditions.

Modulation of LPA signaling has the potential to be of therapeutic value in periodontal disease (PD) and conditions affecting the oral system (reviewed in [16]). Given the solid evidence that LPA regulates salivary gland development, physiology, and is present in saliva, we hypothesized the following in a model of PD using *P. gingivalis*–infected mice: (a) the LPA species found in normal human saliva would be present in mouse saliva, and pharmacologic elevation of one or more of these species would also occur after they developed PD; (b) the adult murine SMG would express LPA4; and (c) immunofluorescence (IF) confocal imaging of LPAR distribution in whole-mount murine salivary gland tissue could be complemented by second harmonic generation (SHG) imaging of collagen to visualize its distribution.

### EXPERIMENTAL RESULTS AND DISCUSSION

Our results show that we successfully established PD using the widely utilized model of *P. gingivalis-*infected C57BL/6 mice **(Fig. 1)**. Specific antibiotic regimens are usually used to suppress the oral microbiome before introducing defined periodontal pathogens to establish a controlled periodontal disease model. The choice to not pre-treat with antibiotics was done to preserve normal flora, given what we now know about the major contribution of the microbiome to homeostasis and to experimental responses and mucosal immunity [36,37]. However, we demonstrated that these mice could indeed develop PD naturally over time, without the usual protocol of antibiotic pre-administration, mimicking how human PD develops.

**Fig. 1.**
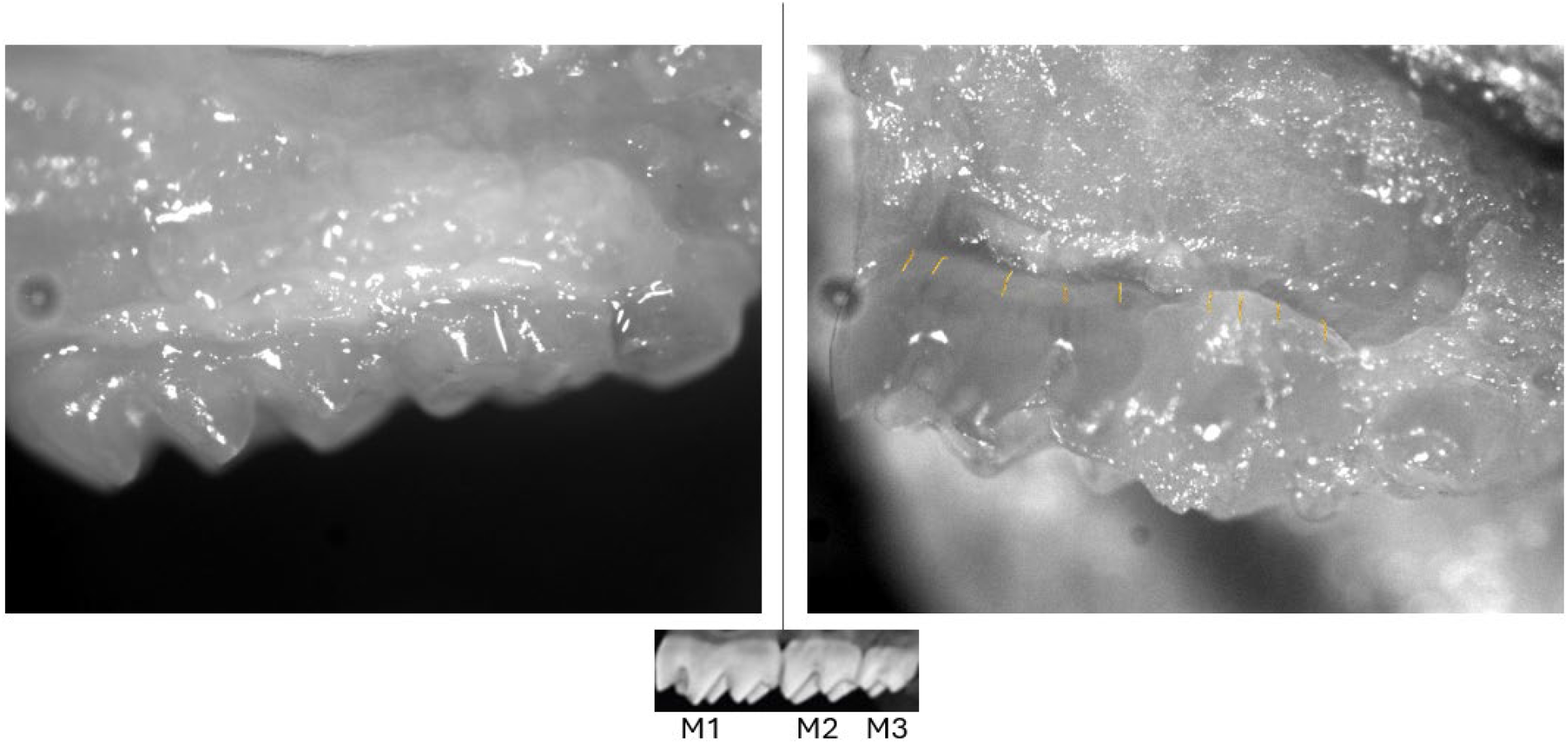
Healthy control and periodontally diseased mice. Representative dissecting microscope images (20x) of maxillae are shown from day 45 post-infection with *P. gingivalis*. Small reference picture below shows positions of molars (M) 1, 2, and 3. Left panel, control mouse showing healthy gingiva adhering tightly to molars M1-M3. Right panel, periodontally diseased mouse, showing gingival recession (recession distance from original location indicated at multiple points with gold lines for M1 and M2). The darkened gingival areas show swelling and bleeding, and M3 also shows some tooth discoloration.

Human mixed saliva contains LPA18:1 > LPA18:0 > LPA16:0 > LPA18:2 (3,4]. Just like we found for PD patients [8], this mouse PD model is also associated with a similar (∼10-fold) salivary LPA species elevation, especially for LPA 16:0 and 18:1 – but only in males (**Fig. 2**). This is interesting to note, as there is known sexual dimorphism in the mouse salivary glands; Mukaibo et al. (2019) [38] undertook “to identify the sex-specific differences in the transcriptional profiles of the three major mouse salivary glands that may have functional consequences.” They reported striking transcriptome differences between male and female glands. They also found a very strong male bias for sex-enriched genes in sublingual glands, but the SMG was notable for the greatest expression of sexually dimorphic genes. Their findings likely explain the salivary LPA sex differences we found in this study.

**Fig. 2.**
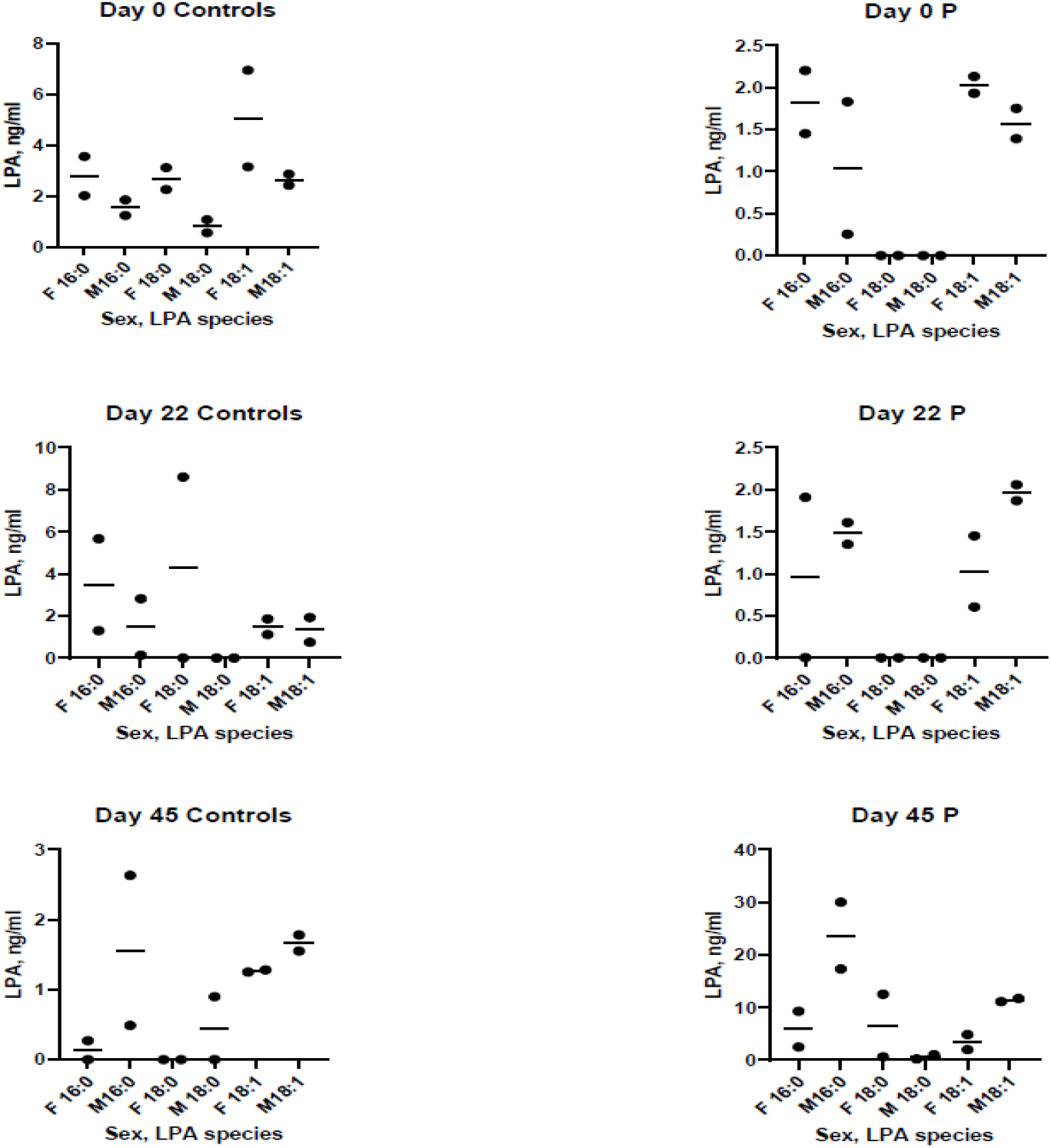
LPA species present in whole saliva from normal control and periodontally diseased mice. Days 0, 22, and 45 post-infection with *P. gingivalis*. Each point represents 2 pooled saliva samples (see “Experimental Results and Discussion” for rationale). For periodontally diseased males only, day 45 shows elevation of LPA 16:0 and LPA 18:1. F= female; M= male; 16:0= LPA 16:0; 18:0= LPA 18:0; 18:1= LPA 18:1; C= Control mice (sham-infected); P= Periodontally diseased mice.

Our sample size calculation from the statistician indicated that for a two-tailed t test with a significance of α = 0.05, a sample size of 7 mice per group would provide 99% power to detect a 20% difference in alveolar bone loss between the *P. gingivalis*-infected mice and the *P. gingivalis*-infected mice with the LPA antagonist we were testing in our grant. So, we rounded to 8 mice per group (8 females and 8 males). This number may be enough animals to satisfy statistical validity but is still a small group.

To the best of our knowledge, no one has measured LPA species in mouse saliva until this study. We were unsure as to whether the scant microliter quantities of saliva collected from individual mice would allow for sensitive LPA species detection in one animal-so the paper points from two mice were pooled and analyzed together and then graphed as one point. As C57BL/6J mice are inbreds, we felt pooling would be justified to make sure we could detect the LPA species. Therefore, due to the small number of mice tested and pooling of samples, the raw data is shown in **Fig. 2** as a scatter plot, but without statistical analyses.

Hwang et al. [39] reported the presence of all currently International Union of Basic and Clinical Pharmacology (IUPHAR)-designated LPARs (LPA1-6) by RT-PCR in the human submandibular gland (HSG) cell line, and elevation of intracellular Ca2+ in response to LPA (presumed to be LPA 18:1 as it is the most used species, since they did not supply that information). However, Lin et al. (2018) [40] used short tandem repeat analysis to analyze HSG from several laboratories, which revealed contamination by the ubiquitously used HeLa (human cervical carcinoma) cell line, revealing “>80% match with HeLa in both the ATCC and Deutsche Sammlung von Mikroorganismen und Zellkulturen (DSMZ) databases.” As genotyping was not reportedly performed during HSG cell line establishment, it is difficult to authenticate if any laboratory is actually using uncontaminated HSG unless individual laboratories have tested their HSG cells.

We show that LPARs can be visualized by confocal IF in ∼ 4 × 2 × 1 mm-thick pieces of mouse SMG (**Fig. 3**), using a whole-mount technique we developed for human gingival tissue and periodontal ligament *in situ* detection of LPARs [41] followed by clearing the tissue. To our knowledge, this is the first report of the presence of LPA4 in the adult mouse SMG. The ability to use intact SMG to study and monitor GPCR expression and distribution represents a useful and substantially time-saving method over having to embed and section the tissue and has the great advantage of providing panoramic spatial information, as it preserves the intact architecture of the gland and allows for enhanced interpretation of biological relationships.

**Fig. 3.**
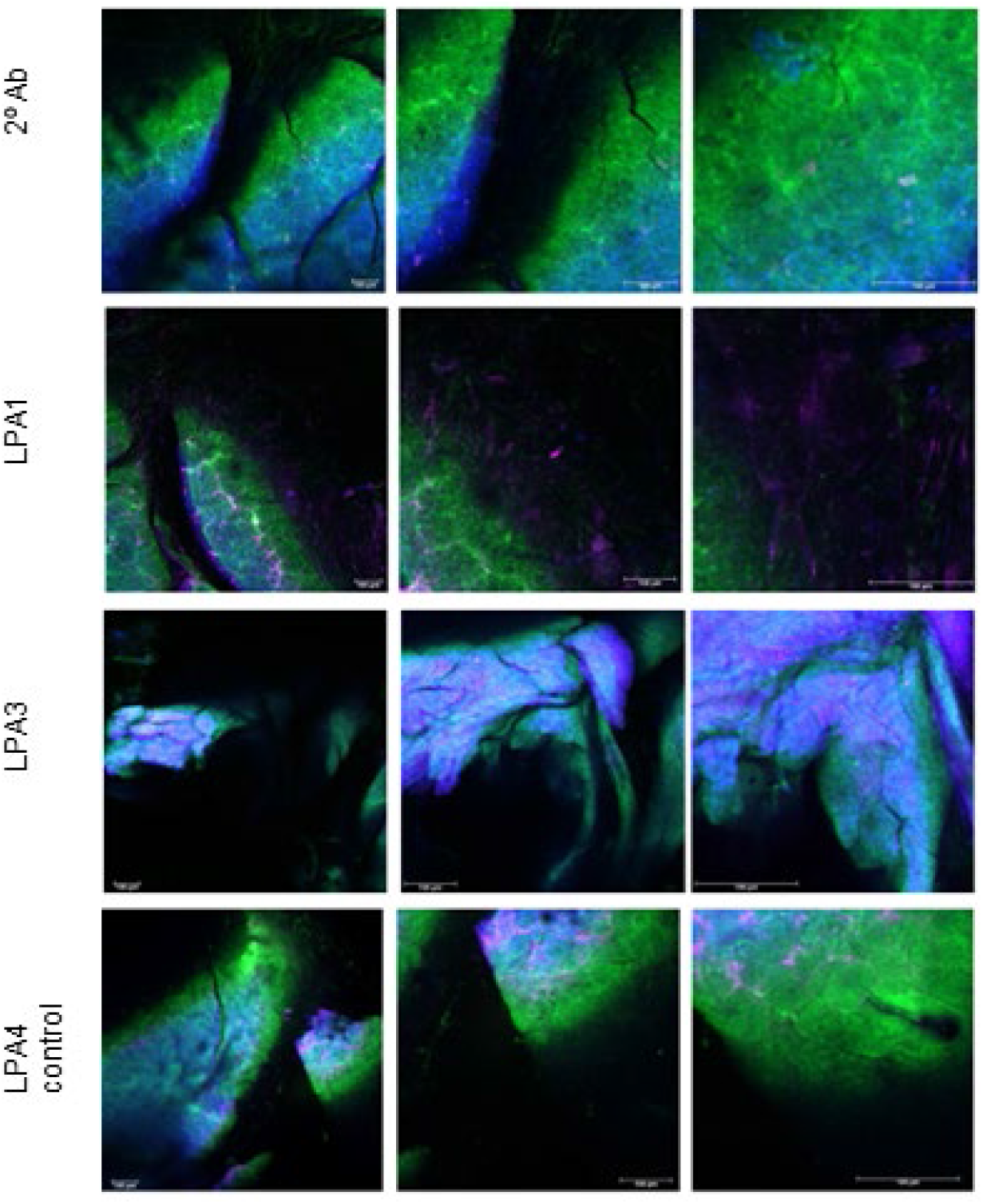
LPA1, LPA3, and LPA4 expression and collagen distribution in adult mouse whole-mount SMG tissue. Representative confocal images of healthy SMG tissue merged with SHG collagen visualization are shown. Left to right: 10x, 20x, and 40x magnification. Note that full field of view for the images is: 10x, ∼ 1.1 mm, 20x, ∼ 0.53 mm and 40x, ∼ 0.28 mm. Scalebar = 100 microns. DAPI nuclear label, blue; LPAR antibody label, magenta. SHG analysis (green, verified by fluorescence lifetime imaging) shows the distribution of collagen fibers. Confocal images are shown as merged signals. For LPA1 (40x) the SHG was obscuring the fluorescent antibody signal -so it was reduced to enhance visibility of the LPA1 label. LPA1 appears distributed in the large ducts and in the gland capsule; LPA3 is mostly distributed in the gland parenchyma, while the LPA4 label appears mostly in the ducts.

Unlike Park et al. (2017) [27] who did not detect expression of LPA4 nor LPA5 in the salivary glands of nonobese diabetic (NOD) mice model of Sjögren’s disease, we were able to detect it in our adult C57BL/6J [and also in SHK1 mice SMG (data not shown)]. Confocal detection with anti-LPAR antibodies showed that LPA1 and LPA4 appear to be mainly expressed by duct cells; LPA1 labeling is also visible on the capsule of the lobule (**Fig. 3**, 10x magnification). LPA3 showed wide expression throughout the SMG.

SHG is a nonlinear optical phenomenon in which two photons of identical wavelength interact within a non-centrosymmetric material (like fibrillar collagen) to create a new photon whose wavelength is precisely half that of the incoming photons. Masedunskas & Weigert (2008) [42] showed that intravital imaging using a combination of two-photon and SHG yielded beautiful images of the rat SMG, showing the outline of cells, internalization of molecules like fluorescently labeled dextran, and visualization of large elastic fibers; but that technique is expensive and requires special equipment.

**Fig. 3** also shows that the concurrent LPAR/SHG analysis of the SMG effectively revealed the architecture of collagen fibers. It was not our intent (as the LPAR detection in whole-mount SMG was a side undergraduate summer project - and not part of the grant) to characterize the cells expressing these LPARs, but to document their presence and overall distribution in the adult mouse SMG- and to provide proof of principle for concurrent LPAR and collagen detection and visualization using whole-mount, clarified SMG tissue.

Being able to visualize the density and organization of collagen fibrils provides a much-needed, complementary method for later studies to use to investigate inflammatory changes in this important ECM molecule in salivary glands relative to the expression of LPARs, which are key in both homeostasis and inflammation.

## CONCLUSIONS

Our findings indicate that we successfully induced PD using the widely adopted model of *P. gingivalis*–infected C57BL/6J mice, which showed very similar ∼10-fold elevation of two of the several predominant salivary LPA species that we found in patients [8]. Notably, we also showed that these mice can naturally develop PD over time without the standard practice of antibiotic pre-treatment, providing a model that more closely reflects the progression of PD in humans.

This pilot study showed that LPA1and LPA4 expression appears to be more restricted to duct cells, while LPA3 is extensively expressed throughout the SMG parenchyma. To our knowledge, this is the first report of LPA4 being expressed in the adult mouse SMG. Concurrent SHG imaging effectively revealed the density and 3-D organization of collagen in the mm-sized, whole-mount mouse SMG fragments. Most PD models use the mouse, but the association / involvement of salivary gland LPARs in PD has not been investigated; defining the expression of salivary gland LPARs in the mouse takes a step in that direction for future investigations.

The presence and distribution of these LPARs in the SMG strongly supports our contention that LPARs are likely to participate in regulating both homeostasis and inflammation in the salivary glands. The presence of LPA4 on the SMG duct cells suggests that this receptor may serve as a regulatory receptor to LPA1 as reported for bone [13]; this remains to be determined in future studies.

## MATERIALS and METHODS

### Periodontal disease induction

All mouse experiments were conducted under the approval, guidelines, and regulations of Creighton University’s Institutional Animal Care and Use Committee (IACUC), Protocol number 1106.1 (all authors complied with ARRIVE guidelines).

*P. gingivalis* strain FDC 381 [(ATCC^®^ BAA1703^™^) was purchased from American Type Culture Collection [(ATCC), Manassas, VA, USA]. All work with *P. gingivalis* was done in an anaerobic hood with an atmosphere of 80% N_2_, 10% H_2_, and 10% CO_2_. All bacterial culture plates and liquid media were equilibrated in the anaerobic hood for at least 12h to remove residual oxygen prior to being inoculated. The bacterial culture from ATCC was streaked onto trypticase soy blood agar plates (TSA II with 5% sheep blood) supplemented with hemin and menadione (Sigma-Aldrich, St. Louis, MO, USA). The plates were incubated for 48-72 h until colonies formed, then the colonies were transferred to Brain Heart Infusion (BHI) broth (Sigma-Aldrich) supplemented with hemin and menadione to grow for the infective cultures needed. The cultures were grown until mid-log growth (OD_660_ 0.5-0.7), then harvested by centrifugation (5,000 × g, inside the anaerobic hood). The pellets were gently resuspended in sterile phosphate-buffered saline (PBS) with 2% carboxymethylcellulose and adjusted to ∼ 10^10^ CFU per100 μl.

Ten-week-old, non-antibiotic-pretreated JAX® C57BL/6J mice (male and female; Jackson Labs, Bar Harbor, Maine, USA) were kept under standard environmental conditions (22–24 °C, 12 h light/dark cycles) with free access to water and standard mouse chow. They were acclimatized for one week post-shipment before experiments were begun in order to eliminate/minimize shipment any stress-related effects.

They were chronically exposed to the live 10^10^ CFU of *P. gingivalis* in 100 μl of PBS/mouse in the 2% carboxymethylcellulose vehicle, or vehicle alone (sham-infected controls) gently applied to all their gingivae using a sterilized ball-tipped steel gavage needle (Thermo Fisher Scientific, Waltham, MA, USA), 5x/week for three weeks. To ensure that their gingivae could be thoroughly, completely, and reproducibly covered with the foul-smelling bacterial suspension, the mice were first anesthetized with isoflurane just until tail drop, then rapidly removed from the anesthetic chamber and treated with the *P. gingivalis* suspension.

### Saliva Collection for LC-MS/MS LPA Species Analysis

Saliva was collected by using a sterile dental paper point [size 0.04, Dia-ISOGT Paper Points (Henry Schein, Melville, NY, USA)] for 5 seconds to soak up ∼2 μl of saliva from each animal’s mouth (back right mandible, near lower molars). Each mouse was sampled on day 0 before infection, on day 22, and then on day 45 before sacrifice (all the mice in the experimental group developed full-blown PD). The collected paper points were immediately frozen at -80 ^o^C until analysis as per [8]. The mice were euthanized by isoflurane inhalation overdose.

### LC-MS/MS LPA Species Analysis

LC-MS/MS analysis of LPAs was performed following the method we previously described in [8] with minor modifications: Chromatographic separation was carried out using a Waters ACQUITY UPLC system (Waters, Milford, MA, USA) coupled to a 4000 Q TRAP® hybrid triple quadrupole/linear ion trap mass spectrometer equipped with an electrospray ionization (ESI) source (Applied Biosystems/MDS Sciex, Foster City, CA, USA). Data acquisition and instrument control were performed using Empower Pro 6.0 and Analyst 1.4.2 software, respectively.

Separation was achieved on a Macherey-Nagel NUCLEODUR® C8 column (125 × 2.0 mm, 5 µm), which was protected by a C18 guard column (Waters, Milford, MA, USA). The mobile phase consisted of solvent A, methanol/water (75:25, v/v) containing 0.5% formic acid and 5 mM ammonium formate, and solvent B, methanol/water/formic acid (99:0.5:0.5, v/v) containing 5 mM ammonium formate. A gradient elution program was used, beginning with 100% solvent A, increasing to 100% solvent B within 1 min, holding at 100% B for 2.5 min, and then returning to the initial conditions. The total run time was 8.5 min. The flow rate was set at 0.5 mL/min, the injection volume was 10 µL, and the column temperature was maintained at 23 °C. Mass spectrometric detection was conducted in negative ion mode using optimized parameters, including an ion spray voltage of −4000 V and a source temperature of 700 °C. Nitrogen was used as both curtain and collision gas. Quantification was performed by monitoring multiple reaction monitoring (MRM) transitions for LPA species in saliva samples.

### Salivary glands

SMG were rapidly dissected from mice on ice after the isoflurane overdose euthanasia and fixed in 10% neutral buffered formalin until analysis.

For confocal imaging, approximately 4 × 2 × 1 mm SMG pieces were cut, de-lipidated in ice-cold 1:1 methanol-acetone for 25 min at -20 C, rehydrated through graded alcohols, and extensively washed in Tris-buffered saline (TBS) with Tween (0.5%), (TBS-T) to prepare for antibody labeling. The SMG pieces were then blocked overnight at room temperature (RT) in TBS-T with 2% gelatin and 0.1% azide.

The SMG pieces were probed with anti-LPAR rabbit polyclonal antibodies (LPA_1_, **#**10005280; LPA_3_, #10004840, both from Cayman Chemical, Ann Arbor, MI, USA) or LPA_4_ polyclonal antibody #LS-C402500-60 (LS Bio, Seattle, WA) overnight at RT in TBS-T with 2% bovine serum albumin (BSA) followed by extensive washes in TBS-T at RT for 6 h with hourly buffer changes. The same sequence was used to label the SMG pieces with goat anti-rabbit secondary fluorescent antibody conjugates CF®-532, -588, or -647 (Biotium, Fremont, CA, USA) for LPAR detection by indirect immunofluorescence (IF), followed by extensive washes in TBS-T at RT for 5 h with hourly buffer changes. DAPI (4′,6-diamidino-2-phenylindole) was used to label nuclei. The SMG pieces were then dehydrated through graded alcohols and cleared by transferring into BABB (1:2 benzyl alcohol/benzyl benzoate). The tissues became completely transparent within minutes using this method. Experiments were repeated a minimum of three times.

## Confocal Imaging

Three-color Images of the SMG fragments were obtained using a Leica TCS SP8 MP confocal microscope. First, the nuclear counter stain (DAPI, 405 nm excitation with detection bandpass of 422-472 nm) and labeled LPARs (633 nm excitation with a detection bandpass of 658-711 nm) were obtained using single-photon confocal imaging. Then, an SHG image of the same field of view was obtained using the femtosecond mode-locked pulse train of a Spectra Physics Mai Tai Ti:S laser (Newport Corporation, Irvine, CA, USA) operating at 920 nm with non-descanned detection by a Super HyD detector with a detection bandpass of 420-500 nm (HQ 460/80, Chroma Technology, Bellows Falls, VT, USA). Fluorescence lifetime measurements performed with a time-correlated single photon counting module (SPC 830, Becker and Hickl, Berlin Germany) confirmed prompt SHG was the most significant emission following 920 nm illumination. 10x images were obtained with a HC PL Fluotar 10x/0.30 objective (Leica).

### Dissecting Microscope Images

Images of the gingivae and mouse molars (M1, M2, and M3) on the mandibles and maxillae were obtained using an Olympus Stereo Microscope SZX12 with camera attachment.

### Data Analysis

Saliva LPA species data were graphed with GraphPad Prism, GraphPad Software Inc. 9.0.0, https://www.graphpad.com.

## Acknowledgements

The authors wish to thank Lillian Calisto B.S. for excellent technical assistance, and Drs. Nancy D. Hanson and Travis J. Bourret for anaerobic culture advice and for providing the anaerobic culture hood. Anthony Stender Ph.D., Manager of The Integrated Biomedical Imaging Facility (IBIF) at Creighton University provided valuable guidance for the confocal imaging.

## Author contributions

Conceptualization, D.R.C, M.G.N., T.M.; Methodology, D.R.C, M.G.N., T.M., D.K., Y.A.

Investigation, D.R.C., D.K., M.G.N., G.R.R., M.E.F.

Writing –Original Draft, D.R.C., D.K. Writing – Review & Editing, D.R.C., D.K., M.G.N, T.M., Y.A. G.R.R, M.E.F.

Visualization, D.R.C., T.M.; Funding Acquisition, D.R.C.

## Data availability statement

This work did not generate publicly archivable numerical datasets. The data, however, are available upon request.

